# Benchmarking Reverse Docking through AlphaFold2 Human Proteome

**DOI:** 10.1101/2023.12.16.572027

**Authors:** Qing Luo, Sheng Wang, Hoi Yeung Li, Liangzhen Zheng, Yuguang Mu, Jingjing Guo

## Abstract

Predicting binding of a small molecule to the human proteome by reverse docking methods, we can predict the target interactions of drug compounds in the human body, as well as further evaluate their potential off-target effects or toxic side effects. In this study, we constructed 11 pipelines to evaluate and benchmark thoroughly the predictive capabilities of these reverse docking pipelines. The pipelines were built using site prediction tools (PointSite and SiteMap) based on the AF2 human proteome, docking programs (Glide and AutoDock Vina), and scoring functions (Glide, Autodock Vina, RTMScore, DeepRMSD, OnionNet-SFCT). The results show that pipeline glide_sfct (PS) exhibited the best target prediction ability and successfully predicted the similar proteins of native targets. This finding provides important clues for understanding the promiscuity between the drug ligand and the whole human proteome. In general, our study has the potential to increase the success rate and reduce the development timeline of drug discovery, thereby saving costs.

## Introduction

Drug discovery is time-consuming and requires a significant amount of funding, with an average duration of over 10 years and a cost of approximately $350 million per drug to reach the market. Furthermore, approximately 90% of drug candidates studied in humans do not successfully progress through clinical trials due to concerns regarding their safety and efficacy.^1,2^ Toxic side effects or ineffectiveness of drugs often stem from the promiscuity of the drug itself. A promiscuous ligand is typically described as a small molecule that exhibits activity against two or more distinct protein targets. ^3^ In the past decades, the one-target, one-drug paradigm has been the dominating drug discovery approach. However, recent advances in experimental and computational studies have shown that small-molecule drugs are rarely selective enough to interact solely with their designated targets. ^4–6^ More than 50% drugs interact with more than five targets, usually resulting in unexpected side effects or toxicity.^5,7–10^ This is likely a conservative view of drug selectivity which is limited by the lack of completeness of drug–target interaction data. After entering the human body, each drug interacts with a series of proteins. Among these interactions, the ones with protein targets relevant to the disease contribute to the main therapeutic effects known as polypharmacology. The most successful long-established antibiotics actually act on multiple targets.^11,12^ However, interactions of drugs with other non-disease-related targets often result in side effects. Drugs may also interact with targets related to other diseases, known as drug repurposing. The essence of all these phenomena can be attributed to the promiscuity of the ligand. Thus, it is critical to identify all potential targets or weaker binders in humans for small molecules to better understand ligand promiscuity. It can anticipate and explain side effects in advance, avoid off-target effects, enhance drug efficacy, and have a higher probability to obtain favorable clinical results.

Protein target prediction, also known as target fishing, helps to identify the potential targets of a query molecule.^13^ As we all know, experimental methods for predicting targets are time-consuming and expensive. Thus, computational target prediction has become an indispensable tool that can rapidly profile a ligand against various macromolecular targets and guide experimental design by only testing fewer proteins. These computational methods can be classified into ligand-based and structure-based methods. ^14^ Ligand-based methods simplify the problem to a similarity-searching problem, only utilizing ligand information to predict targets indirectly.^15^ Compared to other methods, structure-based methods, such as reverse docking, have significant advantages. They utilize the three-dimensional structure of proteins and information about ligand binding pockets to predict the binding mode and strength of a ligand.

Reverse virtual screening has increasingly become a key approach for polypharmacology, drug repurposing, and the identification of new drug targets.^16^ With the rapid growth of available experimental protein structures and the development of reliable protein structure prediction tools,^17,18^ the accuracy of reverse docking to predict the protein-ligand interaction could be enhanced. For example, Cai et al. successfully identified anti-Helicobacter pylori agent targets of an active natural product and its derivative compound using a reverse docking approach. The result was confirmed with the X-ray crystallography.^19^ In a study conducted in 2012, Eric and colleagues employed Tarfisdock to perform reverse docking based on a set of protein targets. Through this approach, they successfully identified potential targets that could elucidate the cytotoxic effects of aryl-aminopyridines and their derivatives.^20^ These successful cases demonstrate the significant contribution of reverse docking to predicting protein targets for small molecules. Reverse docking has emerged as a valuable tool in small-molecule target predictions.

Despite significant advances in reverse docking, this method still faces practical limitations. For instance, there is limited availability of three-dimensional target structures, difficulties in determining the binding sites for small molecules and side-chain orientations, and challenges in selecting appropriate docking programs and scoring functions. In addition, the scoring functions of current docking programs are primarily designed for docking or virtual screening of small molecules, and there are few specifically optimized scoring functions for reverse docking^6^. Interestingly, in large and ultra-large ligand screening practices, current scoring functions suffer from very high false positive rates^21^. Therefore, the same or worse situations would be anticipated in reverse screening tasks.

Here, to comprehensively evaluate the current state of reverse docking for target fishing, we systematically evaluated the impact of various factors using AlphaFold2 predicted structures at the human proteome level (Figure 1). Several factors including binding sites as well as pocket residue flexibilities, docking/sampling methods, and scoring functions may affect the pipeline performance. Our findings provide key insights into the current capabilities and future opportunities for target fishing and drug discovery powered by AI-predicted protein structures. Moreover, the optimal protocol for reverse docking could be selected as a proteome-level target fishing platform, which can be used to discover potential drug targets, evaluate side effects and off-target effects, and accelerate the discovery and development of drugs, including drug repositioning and multi-target or polypharmacological ones.

**Figure 1:**
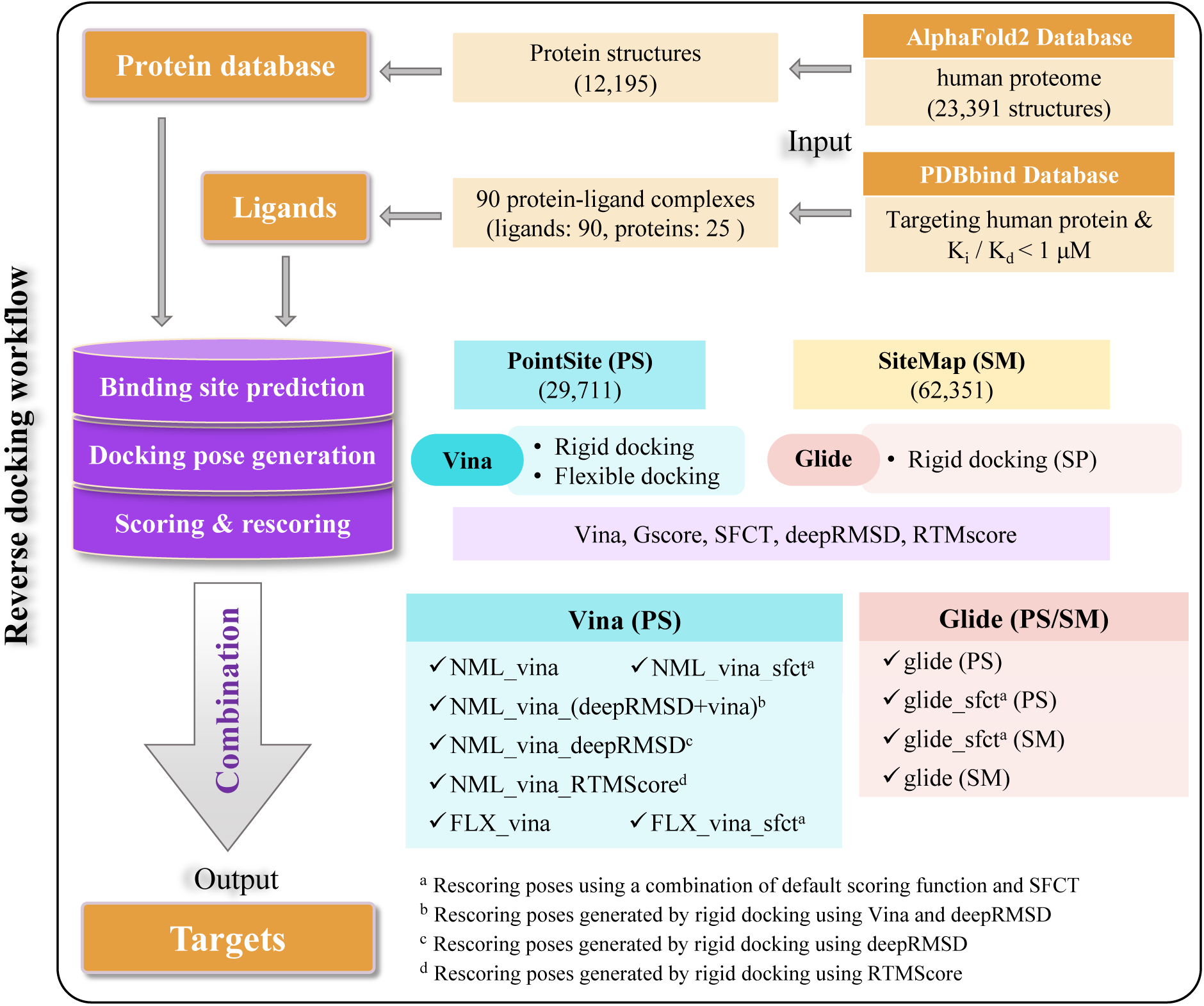
The workflow of reverse docking pipelines in the present study.

## Materials and Methods

### Benchmark Datasets

#### Benchmark Dataset of the Human Proteome

The structures of the proteins in human proteome were downloaded from the AlphaFold Protein Structure Database (https://alphafold.ebi.ac.uk/). ^22^ The total number of proteins is 23,391, and each protein has a unique Uniprot ID. Low-quality structures, such as disordered protein structures or those lacking well-defined ligand binding pockets, were removed. After this filtration, 14,424 protein structures retained (with individual structure sequence lengths of no more than 1,500 amino acids), representing a total of 12,195 actual proteins.

#### Benchmark Dataset of Ligands

The unique ligand dataset for benchmarking was adopted from the PDBbind Core set, version 2016.^23^ It consists of 91 small molecules in complex with 25 human protein targets, where the affinity values (*K_i_*, *K_d_* or *IC*_50_) for these complexes are below 1 uM. The structures of the 91 complexes and the SDF files of ligands were collected from the PDBbind database. By comparing the predicted protein structure from AlphaFold2 (AF2) with the experimental structure, we observed that the native structure of protein Q15370 (PDB ID 4w9h^24^) consists of three domains. However, the AF2 prediction only identified one domain, which does not correspond to the domain involved in binding the small molecule 3JF. Consequently, 90 ligands were selected for the subsequent calculations.

#### Binding Site Determination

The docking-site centers were determined using two distinct algorithmic approaches, SiteMap (SM)^25,26^ or PointSite (PS).^27^ SiteMap is geometry-based and employs a scoring function called SiteScore, which assesses and ranks the suitability of different sites for ligand binding by searching non-protein grid point clusters. Here, with the default setting, we retained a maximum of the top five potential sites for each protein based on the SiteScore ranking.

PointSite is a template-free ligand-binding site prediction algorithm based on a point cloud deep learning model that can provide the probability of each atom in a protein for ligand binding. The pocket region of the protein is defined as the area around atoms with high probabilities. Unlike other geometry-based algorithms (such as Fpocket^28^), PointSite is a protein-centric prediction method that has shown good performance on multiple test sets. In this paper, we employed a method to determine the pocket location based on protein atoms. The specific steps were as follows. PointSite was used to predict the atoms forming the binding sites by assigning scores/probability to each atom. Atoms with binding site probability larger than 0.7 were assigned as binding site atoms. If one binding-site atom is larger than 10 Å away from all found centers of the binding pockets, a new binding pocket will be assigned whose center is the position of this binding site atom. New binding site atoms will be added to the binding site if the distance between the atom and the center of the binding pocket is less than 10 Å. The center of the binding pocket was updated by averaging all constituent binding site atoms. The center position of all found binding pockets was taken as the center for docking. The Uniprot IDs of the proteins and the predicted binding sites by PointSite have been deposited at https://github.com/molu851-luo/Reverse-docking-benchmark.

### Molecular Docking and Rescoring

#### Docking with Autodock Vina

Both normal and flexible docking experiments were conducted using Autodock Vina, a versatile docking software widely utilized in the field. The former involved docking a flexible ligand to a rigid protein, while the latter entailed docking a flexible ligand to a flexible protein. The docking box centers were determined by PS, and the docking box size was 15 Å and the exhaustiveness was 32. For Vina-based flexible docking, the flexible residues were chosen based on the distance between the side-chain atoms and the center of the binding pocket. Three residues with the shortest distance to the binding pocket center were chosen as the flexible residues. Gly, Ala, and Pro residues were not considered flexible residues.

#### Docking with Glide in Maestro

The proteins were prepared using Rosetta, and the Schrodinger LigPrep program^29^ (Schrödinger Release 2023-4: LigPrep, Schrödinger, LLC, New York, NY, 2023.) was utilized for the preparation of ligands with the OPLS4 force field.^30^ The docking box center was determined using two site prediction methods: an internal tool in Maestro, SiteMap, and an external tool, PS. For the reverse docking task, the Standard Precision (SP) docking mode of Glide^31–33^ in Maestro was utilized, employing the default docking parameters. The entire reverse-docking process was executed using the XDOCK script, ensuring a streamlined and efficient workflow. XDOCK is a reverse global docking tool based on the Schrödinger suite of software (https://github.com/Wang-Lin-boop/XDOCK/blob/main/XDOCK).

#### Rescoring Docking Poses of Ligands with Scoring Functions Beyond the Built-in Ones

In addition to the default scoring functions in AutoDock Vina (Vina score) and Glide (Glide-SP Score), alternative methods have also been considered for rescoring the ligand docking poses, including some machine learning-based or deep learning-based scoring methods, such as OnionNet-SFCT (SFCT),^34^ RTMscore^35^ and DeepRMSD.^36^ In this case, SFCT is a correction term used to combine with other existing traditional scoring functions. The combination of Vina with a weight of 0.8.^34^ Here, in the combination of scoring functions Glide-SP and SFCT (with a weight of 0.8). Through a step-wise increment in the proportion of scoring function SFCT from 0.1 to 0.9 at intervals of 0.1, we observed that the optimal performance was attained when its weight reached 0.8, as shown in Figure S1. This finding suggests that SFCT significantly contributed to the achieved results. So, when comparing the performance of the Glide-SP and SFCT combinations with other pipelines, setting SFCT at 0.8 consistently yielded superior results. For each scoring approach, only the best docking pose of a particular protein was kept for data analysis.

In summary, we considered the distinct combination of Glide-SP/AutoDock Vina docking poses, PointSite/SiteMap site prediction methods, and other scoring functions, such as RTMscore, deepRMSD, the combination of SFCT with Glide-SP/AutoDock Vina (Figure 1).

### Assessment Methods

#### Protein Classification

Due to the fact that any scoring function used in molecular docking cannot give reliable predictions for all protein families, the tested proteins were clustered into different protein families, and the prediction accuracy for individual protein families was assessed. In order to explore docking programs and scoring functions’ performance among different protein families, proteins in the test set were clustered into several protein families utilizing the Structural Classification of Proteins extended (SCOPe) database.^37^ As a result, 90 true targets were classified into 15 protein families.

#### Evaluation of Distinct Reverse Docking Protocols

The success rate (SR) is a metric that measures the ability to correctly identify the true target from the top N rankings. The following equation quantitatively defines it:

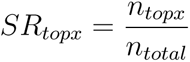

here, the *n_topx_* is the number of true targets ranked in top x. The *n_total_* is the total number of true targets, and the predicted site is in the right pocket. For each ligand, the top 100 ranked proteins are considered as predicted targets. Proteins that are ranked in the top 100 and correctly identified with pockets are considered True Positives (TP), while those that are not correctly identified with pockets are considered False Positives (FP).

The conformation-searching ability and scoring function of current docking programs are not precise enough to reproduce the experimental conformation of all ligands. To focus on the scoring effects, we considered ligands with a small root mean square deviation (RMSD) between their docking and experimental conformations to evaluate the target prediction effect of the 11 reverse docking pipelines.

## Results

### 1. Distribution of the Protein and Ligand Test Datasets

Classifying proteins based on their folds enables a detailed and comprehensive understanding of their structural and evolutionary relationships. It is important to note that the performance of docking tools and scoring functions may vary across different types of protein targets.^38,39^ Therefore, to account for these variations, we utilized SCOPe (Structural Classification of Proteins - extended), a hierarchical classification system, to classify the proteins in our test set. The classification allows us to better understand the results within specific protein fold categories. As a result, the 25 targets were classified into 14 protein families belonging to five main classes (Figure 2A): a, all alpha proteins; b, all beta proteins; c, alpha and beta proteins (a/b); d, alpha and beta proteins (a+b); g, small proteins. However, our test set does not include proteins classified under the main classes e (multi-domain proteins) and f (membrane and cell surface proteins and peptides) in SCOPe. In detail, the target dataset includes a.123.1.1 (Nuclear receptor ligand-binding domain, 1 protein), b.47.1.2 (Eukaryotic proteases, 2 proteins), b.50.1.2 (Pepsin-like, 1 protein), b.74.1.1 (Carbonic anhydrase, 1 protein), c.45.1.2 (Higher-molecular-weight phosphotyrosine protein phosphatases, 1 protein), d.92.1.11 (Matrix metalloproteases, catalytic domain, 1 protein), d.122.1.1 (Heat shock protein 90, HSP90, N-terminal domain, 1 protein), d.144.1.7 (Protein kinases, catalytic subunit, 7 proteins) and g.3.11.1(EGF-type module, 1 protein). Additionally, there are automated matches in the following protein families: a.29.2.0 (1 protein), a.211.1.0 (1 protein), b.47.1.0 (1 protein), c.94.1.0 (1 protein), and d.144.1.0 (2 proteins).

**Figure 2:**
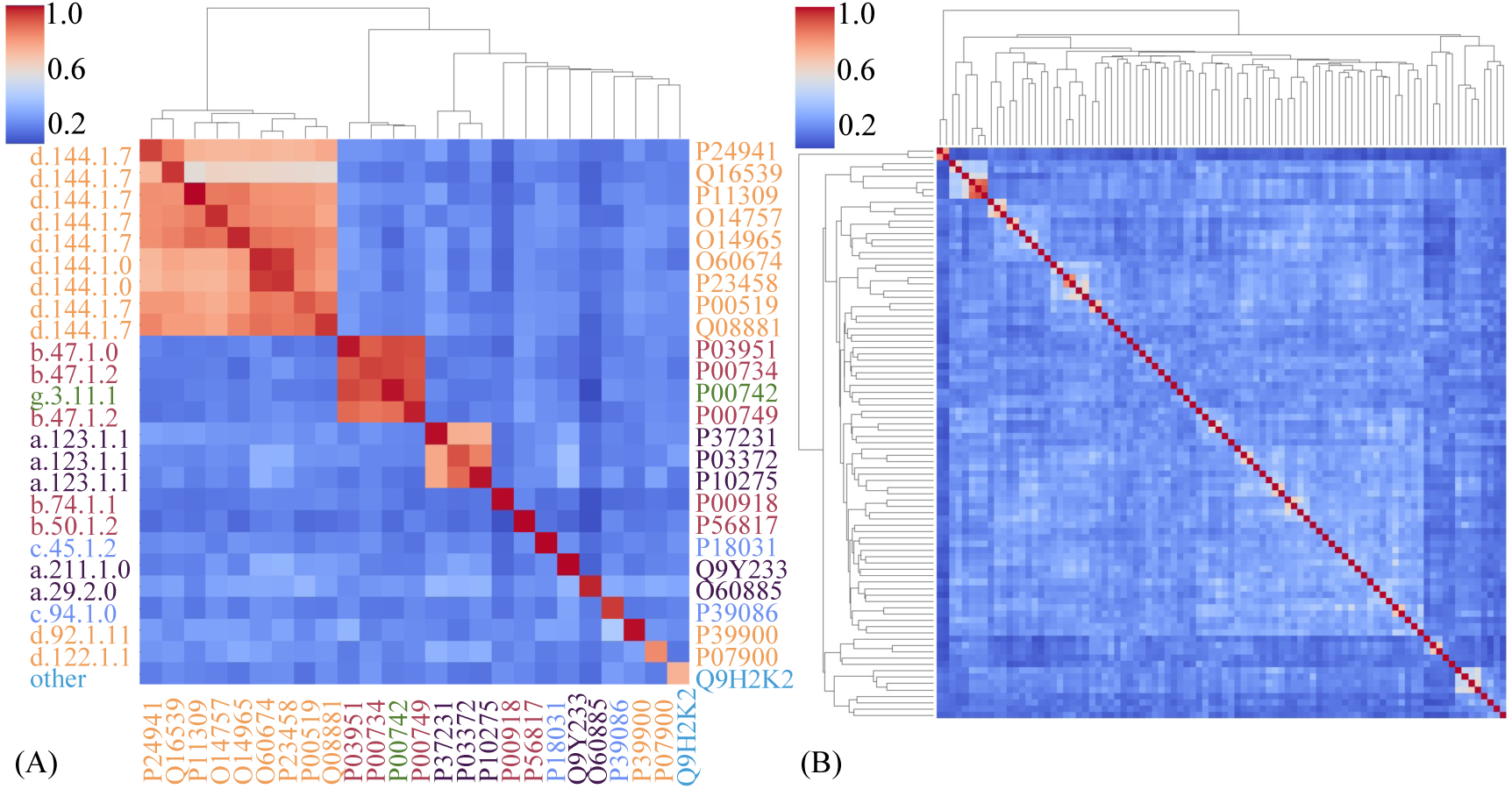
Hierarchical cluster analysis heatmap: The result of cluster analysis of the similarity for (A) small molecules and (B) proteins. Proteins from the same protein family are displayed in the chart using the same color. There are 25 proteins belonging to 15 different families, categorized according to Scope.

Then, to examine the distribution of proteins and small molecules in the test set, we calculated their similarity (Figure 2A). First, DeepAlign^40^ was used to calculate the similarity (TM-score^41^) between the experimental structure of a specific target and the entire set of AF2-predicted human protein structures. ^22^ Among the multiple experimental structures of a protein, the one with the lowest resolution was selected for comparison. In our test set, three protein groups exhibit structural similarity among the 25 proteins. The first group consists of the following proteins: Q16539, P24941, O14757, O14965, P11309, O60674, P23458, P00519, and Q08881. All of these proteins belong to the d.144.1.0 and d.144.1.7 families. The second group comprises P00749, P00734, P00742, P03951. Lastly, the third group consists of P37231, P03372, and P10275. The remaining proteins show relatively low structural similarity to each other. Specifically, the experimental structure of Q9H2K2 exhibits poor similarity to the AF2 structure, with a similarity score of only TM-score=0.743. Based on an examination of the experimental structure of Q9H2K2 and the AF2 structure, it has been observed that there is a significant difference in the pocket region where the small molecule AJ6 binds. This discrepancy is primarily due to the lower quality of the AF2 predicted structure. Consequently, the identification of the native target Q9H2K2 of AJ6 becomes challenging due to these structural variations.

Next, we used the Morgan fingerprint in RDKit to calculate the similarity matrix of the molecules and assessed their similarity using the Tanimoto similarity score. ^42^ As Figure 2B shows, the majority of small molecules have similarity scores concentrated in the lower range, below 0.4 among the 90 small molecules. However, 25 pairs of small molecules have a similarity greater than 0.5 (Figure S2). Among them, 22 pairs share the same protein target. For example, among the five ligands (15T, 1J5, 1J6, 0NT, and 1Q4) targeting P23458, three of them (15T, 1J6, and 1J5) have a similarity greater than 0.8. In addition, ligands targeting the same protein family might also be similar. For example, JAK and 0NV target O60674 which belongs to the same protein family (non-receptor tyrosine-protein kinase, d.144.1.0) with another target protein (Uniprot ID P23458). Interestingly, the similarity between JAK and 1J5/1J6 is about 0.5 while the similarity between 0NV and 0NT is 0.56. However, a pair of molecules, DFL and WST, is not in complex with the same protein or the same protein family. They are ligands of two unrelated proteins.

In summary, we selected strong binding complexes (where only human proteins were considered) from the PDBbind core set as the test set for reverse docking. ^23^ Regarding similarity, except for the low-quality AF2-predicted structure of Q9H2K2, the experimental and AF2-predicted structures exhibited a high structure similarity with TM-scores larger than 0.8. This shows that the structural quality of the AF2 predictions is of high quality. Similar small molecules are consistently bound to the same protein or protein family in our test set. This observation reflects the general principle that similar small molecules tend to bind to structurally similar proteins. ^43–45^ Despite this, there is one exception where the DFL and WST molecules are similar but bind to two unrelated proteins. These examples highlight the promiscuity of interactions between small molecules and proteins. Most of the small molecules in our test set exhibit low similarity, showcasing diverse structures. This diversity is advantageous for our benchmarking purposes as it enables us to assess the performance of our algorithm across a wide variety of compounds and obtain robust conclusions.

### 2. The Performance of Binding-Site Prediction

In the present work, two ligand-binding site prediction tools, PointSite and SiteMap, have been employed in combination with Glide docking to evaluate the importance of site prediction reliably and accuracy in reverse docking. Our results showed that the performance of the two prediction tools is slightly different (Figure 3). For the 25 predicted protein structures, PointSite predicted a total of 208 binding sites, with 76 of them correct. Successful predictions were made for 71 structures and 21 proteins, resulting in a success rate of 84.0%. On the other hand, SiteMap predicted a total of 452 binding sites, with 70 of them being correct. Successful predictions were made for 70 structures and 22 proteins, resulting in a success rate of 88.0%. In terms of individual proteins, SiteMap has a slightly higher success rate. Although both methods achieved high hit rates, 84.0% for PointSite and 88.0% for SM, there were a few proteins in each method where the original binding sites were not accurately predicted.

**Figure 3:**
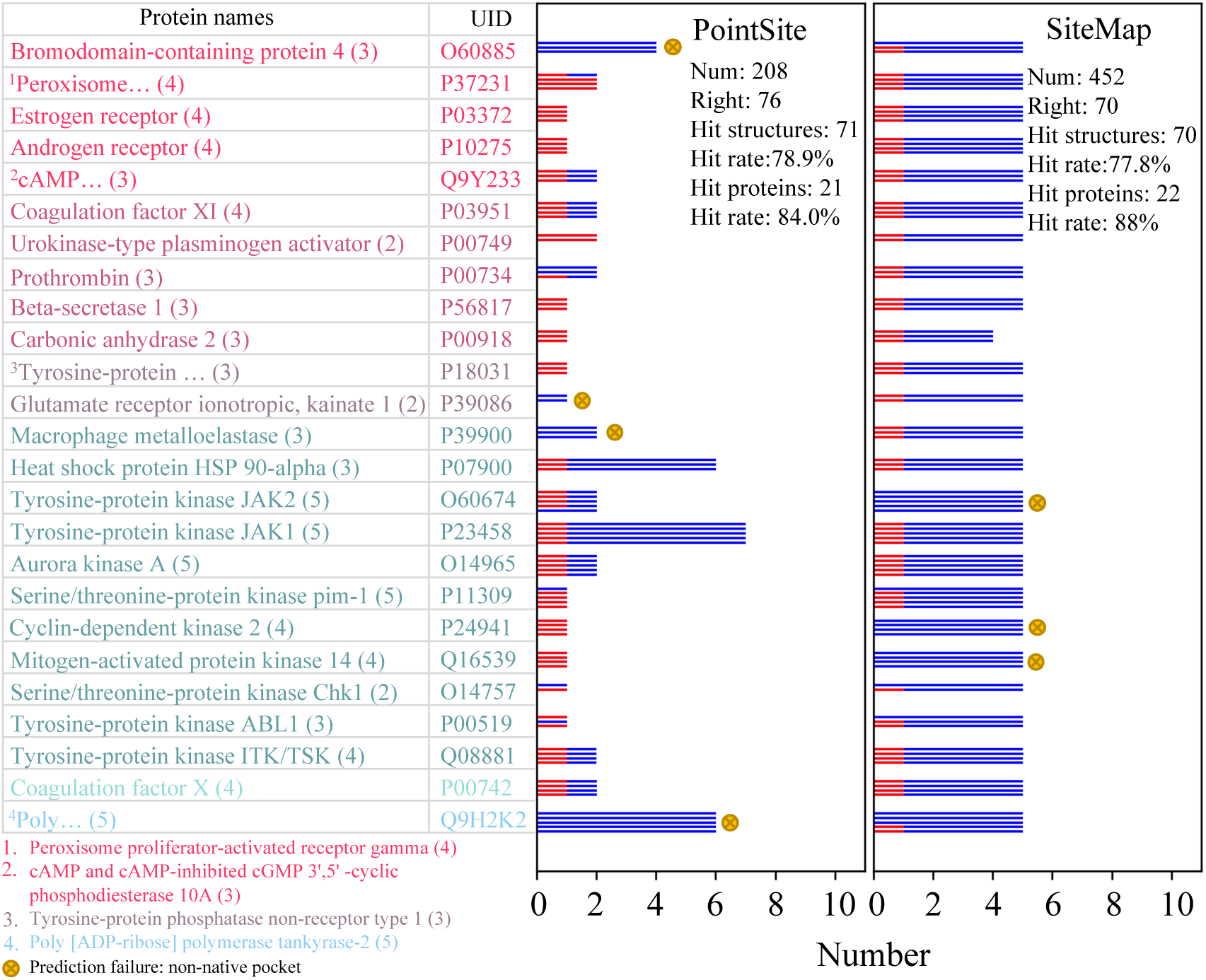
The overview of ligand binding site prediction by SiteMap and PointSite. The red and blue line indicates the number of predicted sites located in the native pocket and non-native pocket, respectively. The total number of sites considered in the docking experiments is 208 and 453 for PointSite and SiteMap, respectively. For each method, there are a few proteins for which the original binding sites were not accurately predicted, and the hit rates for PointSite and SiteMap are 84.0% and 88.0%, respectively.

However, PointSite predicts a lower total number of binding sites in our work. 32 ligands were predicted to have a single binding site that aligns perfectly with the corresponding experimental structure. Furthermore, five ligands were predicted to have two binding sites, and both predictions were successful. The successful site percentage for PointSite is 36.54%. In SM, except for carbonic anhydrase 2 (Uniprot ID P00918), the remaining proteins have 5 sites, with at most one successful prediction, resulting in a successful site percentage of 15.49%. Although the predicted binding sites for the proteins are similar, PointSite has a higher percentage of successful sites, making it easier for the scoring function to select correctly predicted pockets. In addition, there are overlapping and complementary aspects between SiteMap and PointSite. As shown in Figure S5, only SiteMap successfully predicted the binding pocket of protein P39086, while only PointSite successfully predicted the binding pocket of protein Q16539. Additionally, both SiteMap and PointSite successfully predicted the binding pocket of protein P03372.

### 3. Overall Performance of Different Reverse Docking Pipelines

For proteins in the human proteome, we performed systematic docking utilizing two representative docking programs: commercial, Glide (version 67011); ^31–33^ academic, AutoDock Vina (version 1.1.2).^46^ The 90 small molecules were docked against the predicted binding sites of each predicted protein structure in the human proteome. For Autodock Vina, protein flexibility was considered in flexible docking (FLX_vina). As illustrated in Figure 4A, whether it was rigid docking or flexible docking, each molecule was successfully docked with the target by Autodock Vina, yielding pose and ranking results. However, after docking with Glide-PS, only 85 compounds could successfully be docked with the original targets, while Glide (SM) yielded docking successfully for 89 ligands. Assuming the top-ranked proteins as potential targets, Figure 4B provides a clear visualization of the performance of all pipelines with various binding-site prediction methods, docking methods, and scoring functions. As can be seen, the performance of different pipelines exhibits significant variations. Here, to give a more realistic guide for reverse docking and further experimental validation of the predicted targets, we hypothesized that the top-100 ranked targets are potential targets for the query small molecule. Based on the results for the number of origin targets in the top 100, the performance of the pipelines follows the following order: glide_sfct (PS) (27.8%) *≈* glide_sfct (SM) (24.4%) > FLX_vina_sfct (15.6%) = NML_vina_sfct (15.6%) > glide (SM) (10.0%) *≈* glide (PS) (8.9%) *≈* deepRMSD_NML_vina (8.9%) > NML_vina (1.1%) *≈* FLX_vina (1.1%) *≈* deepRMSD (0.0%).

**Figure 4:**
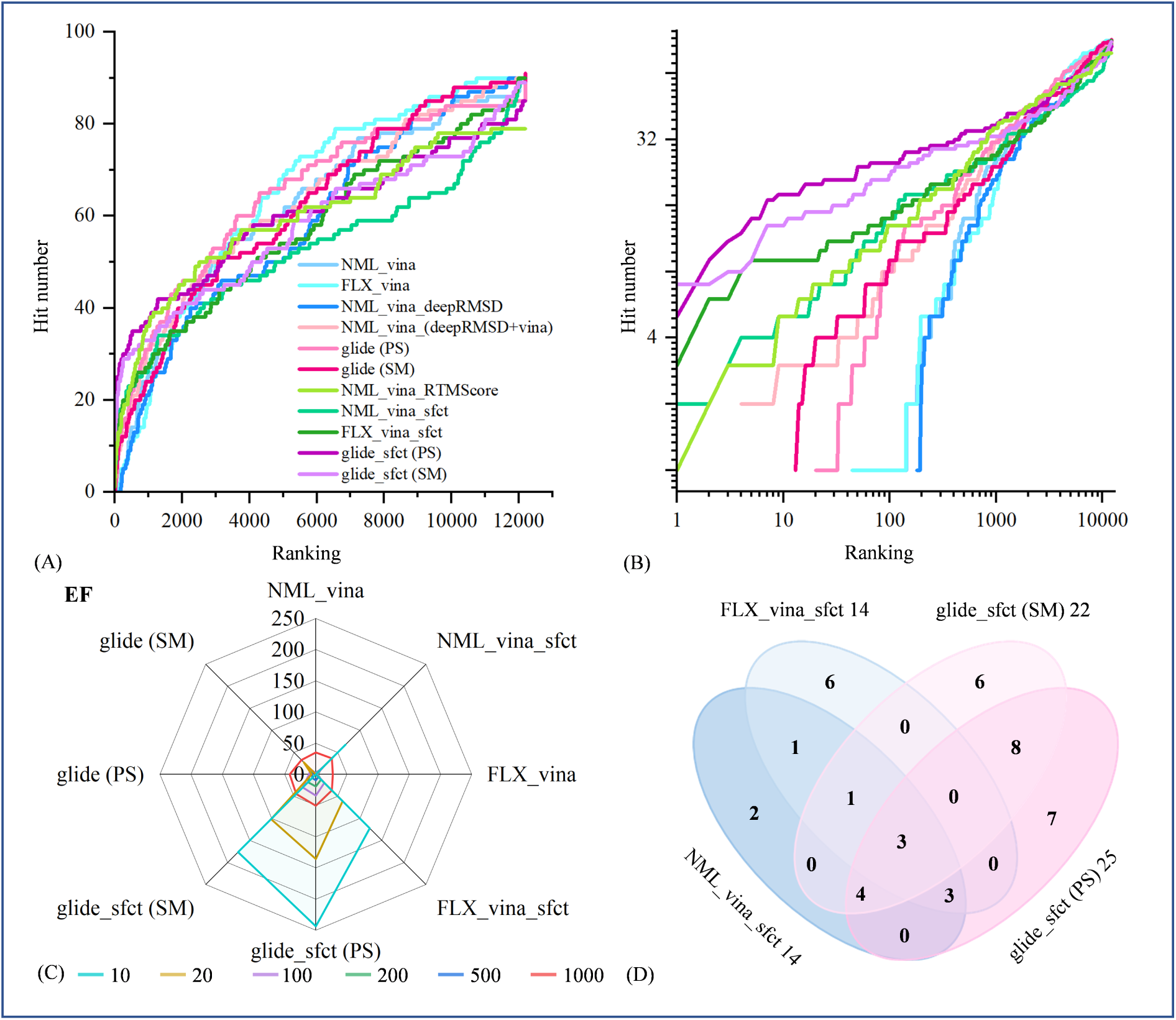
Comprehensive performance assessment of distinct reverse-docking pipelines. (A–B) The plot of hit number as a function of ranking. (C) Radar chart of the performance according to EF. (D) Venn diagram depicting the commonalities and differences among the hits in the top 100 ranking of four representative pipelines, NML_vina_sfct, FLX_vina_sfct, glide_sfct (PS), and glide_sfct (SM).

Furthermore, these pipelines can be categorized into four groups based on performance. The group performed the worst for the top 100 predictions, including NML_vina, FLX_vina, and deepRMSD. Among them, NML_vina and FLX_vina only successfully identified one original target each, while deepRMSD did not make any correct predictions. The second group includes glide (PS), glide (SM), and deepRMSD_NML_vina. Among the top 100, the respective hit numbers for these three pipelines are 8, 9, and 8. Compared to Vina, Glide seems to exhibit superior target prediction ability. However, when Vina was combined with deepRMSD, the performance was close to and even better than Glide. For example, three of the top ten targets are correct in the results of deepRMSD_NML_vina. The third group includes NML_vina_sfct and FLX_vina_sfct. In the top 10 predictions, NML_vina_sfct recognized 5 correct targets, while FLX_vina_sfct had 9. The latter showed a 44.4% improvement over the former one. However, both pipelines successfully predicted 14 targets in the top 100. The fourth group exhibited the best performance, including glide_sfct (PS) and glide_sfct (SM). Surprisingly, Glide_sfct (SM) successfully predicted 13 targets within the top 10 rankings, and Glide_sfct (PS) achieved 18 successful predictions. Overall, glide_sfct (SM) successfully predicted 22 targets, while glide_sfct (PS) performed even better, successfully predicting 25 targets.

It should be noted that before combining with Vina, deepRMSD’s performance was equally poor compared to that of Vina. The number of hits in the top 100 predictions for Vina, both rigid docking (NML_vina) and flexible docking (FLX_vina), was 1. However, when NML_vina was combined with deepRMSD, the number increased to 8, surpassing the performance of Vina and deepRMSD alone. This improvement moved it from the fourth tier to the third tier and produced results comparable to Gscore. After combining with SFCT, the prediction abilities of Vina and Gscore were greatly improved. The glide_sfct (PS) demonstrated the best performance, with an EF value of 33.9. Its performance was outstanding in the top 10 predictions, with an EF value reaching 243.9. Additionally, before the correction by SFCT, glide (SM) performed better than glide (PS). However, the glide (PS) outperformed the glide (SM) in prediction accuracy after SFCT correction. Overall, the integration of the SFCT correction term effectively improved the performance of both Vina and Glide in reverse docking, emphasizing the importance of post-docking rescoring to enhance the effectiveness of reverse docking.

To compare the similarities and differences among the predicted targets of NML_vina_sfct, FLX_vina_sfct, glide_sfct (PS), and glide_sfct (SM), we can refer to the Venn diagram (Figure 4D). Different methods may not achieve a high level of protein structural alignment independently. This indicates that they can complement each other to some extent. Based on the specific screening results shown in FigureS6, we can conclude that the combination SFCT with Glide demonstrates stronger robustness in handling small molecules compared to Vina. When targeting the same protein, Glide often achieves stable and successful predictions of the binding sites for several small molecules. Our reverse docking experiment suggests that using as many as possible ligands in reverse docking is worthwhile in finding unknown drug targets or unexpected mode-of-action even though it costs high computation costs.

### 4. Reverse Docking Performance on Each Test Target

Many proteins in the human body have homologous proteins that perform different physiological functions. ^47,48^ For example, there are various steroid hormone receptors in the human body with a remarkably high level of structural similarity. This creates a situation where small-molecule drugs may display diverse activities when interacting with these homologous proteins.^49,50^ Consequently, this can lead to varying degrees of side effects caused by the drug molecules. However, in specific drug development contexts, the small-molecule drug needs to exhibit a suitable affinity spectrum across a wide range of homologous proteins, and this places extremely high demands on algorithms that predict the affinity and specificity between small molecules and proteins.^51–53^

Proteins exhibit structural similarities and, in many cases, share a common evolutionary origin. According to the structural similarity between the highest-resolution experimental structure of each test target and all AF2-predicted structures, it is evident that the distributions of each evaluated target class are distinct (Figure S4). Some test proteins exhibit a high degree of structural similarity to other proteins in the human proteome, whereas others do not show significant levels of similarity. In our test set, class d (alpha and beta proteins (a+b)) proteins account for the highest proportion. Compared to other protein families, this particular family also shows a relatively high number of similar proteins in the human proteome (Figure S4). Hence, the interaction network between these targets and small molecules tends to be more complex, and proteins similar to the original target have a high probability of being identified through reverse docking.

To further evaluate the ability of different scoring functions to recognize homologous proteins in our reverse-docking benchmark, we analyzed the distribution of similarity between the highest-resolution experimental structure of the original target for each small molecule and the top 100 predicted targets. Receptors derived from the same UniProt ID were consolidated into a single entity. Figure 5 shows that the uncorrected pipelines of Vina and Glide rarely identified targets with high protein similarity. However, after correction, the situation is completely different, and many proteins similar to the original target were successfully identified, especially for targets such as O60674, P23458, and P24941. Interestingly, after SFCT correction, although some targets were not successfully predicted, there were often proteins with high similarity that were predicted, such as P10275, P03951, O14757, and Q9H2K2. Among these pipelines, glide_sfct (PS) performed the best. Out of the 25 proteins with a similar structure to the original target, analogous proteins were successfully predicted for 23 of them, resulting in an impressive success rate of 92%. Among the 90 small molecules, 70 of them had results that included proteins with a similarity score of over 0.6 to the target, resulting in a success rate of 77.78%. In conclusion, the scoring function corrected by SFCT demonstrates greater robustness in predicting targets.

**Figure 5:**
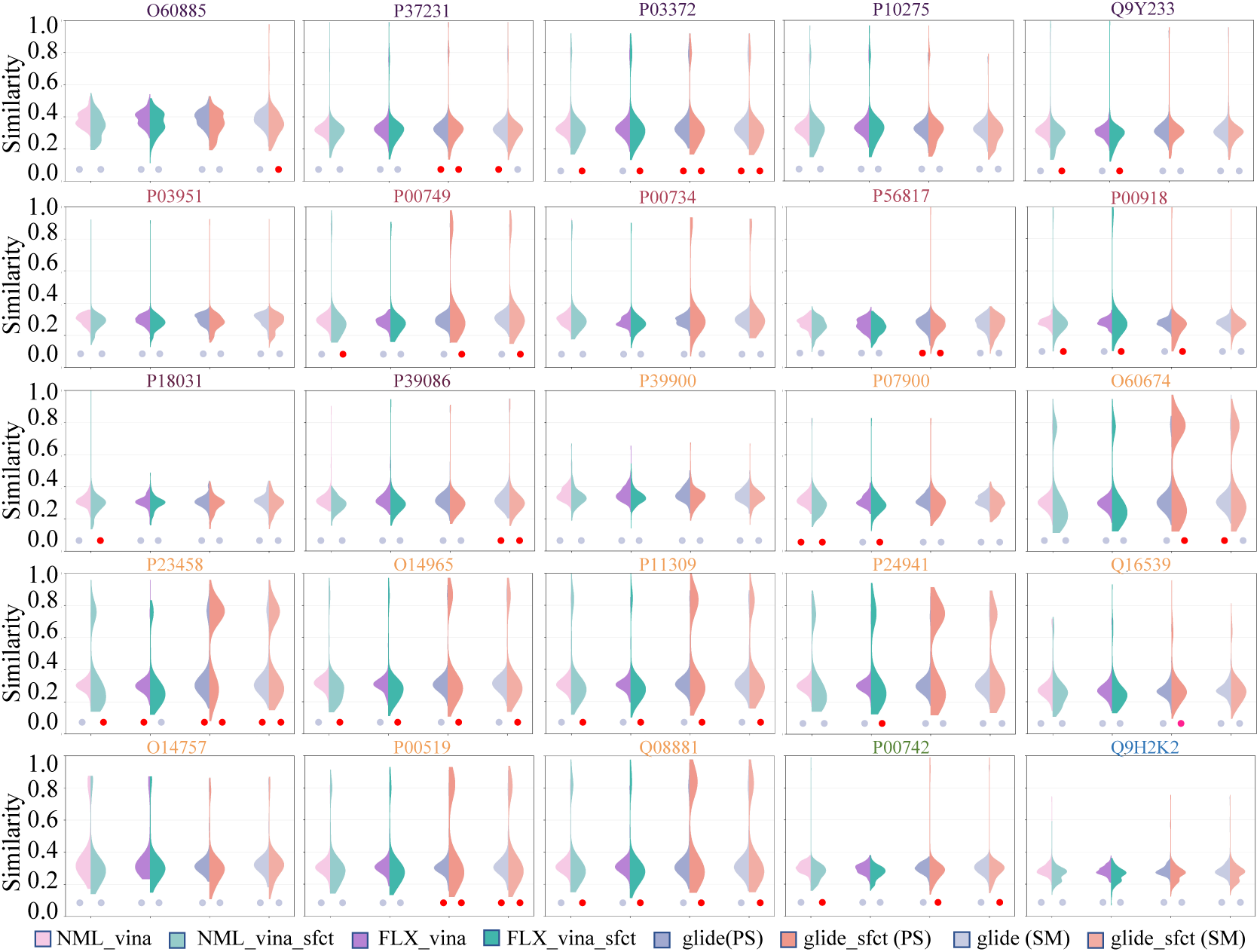
The distribution of similarity between each testing protein and the top 100 hits. In each subplot, red and gray dots represent the successfully and unsuccessfully captured original target by the pipeline, respectively. Each subplot consists of four violins, representing the comparison before and after SFCT correction. Different protein families are represented by UniProt ID in different colors, starting from class a and arranged in sequential order. The red dot represents successful predictions of the original targets and the gray dot represents failed predictions.

Different protein families exhibit varying performance with the same scoring function. The class d has the highest number of proteins, and many proteins, including those with high similarity, are successfully predicted. The four pipelines combined with SFCT show good prediction results in this family. However, the remaining families perform poorly because they have fewer proteins with high similarity. If the quality of the AF2 pocket is not high, it is challenging to predict targets in these families.

## Discussion

### 1. Reverse Docking Accuracy in the Global Queue Scenario

In our work, we used protein structures predicted by AF2 and predicted the binding pockets of small molecules using tools such as PointSite or SiteMap. Therefore, although some native targets are ranked within the top 100, this does not necessarily mean that the small molecules are located in the default pockets. It would be interesting to know (1) whether the ligand binds in the anticipated binding regions (when aligned to the experimental structures) in its destined target protein, and (2) how these accuracies would affect the overall reverse-docking performance.

Here, the docking site (or binding pocket) is thought to be correctly predicted, when the RMSD value of the docked ligands with respect to the corresponding native ligand conformation in the original PDB structure is below 10.0 Å. We consider such reverse docking successful. As shown in Figures 3 and 6, PointSite successfully predicted the binding sites of 76 small molecules, and then the NML_vina_sfct pipeline performed the best with 53 small molecules successfully reverse-docked into the original binding pocket. Additionally, SiteMap successfully predicted the binding sites of 70 small molecules, with the glide (SM) pipeline (47 successfully reverse-docked ligands) performing better than the glide_sfct (PS) one. Therefore, the success rate of reverse docking is quite good. Autodock Vina and Glide perform similarly and do not show significant differences. However, the pipelines involving SiteMap (glide_sfct and glide_sfct (SM)) identified fewer correct binding regions than those of PointSite (Figure 6). There may be two reasons for this: first, Pointsite has a higher success rate in predicting binding sites for small molecules compared to Sitemap; second, SiteMap predicts a larger number of binding sites, making it more challenging to rank the correct site as the best one.

**Figure 6:**
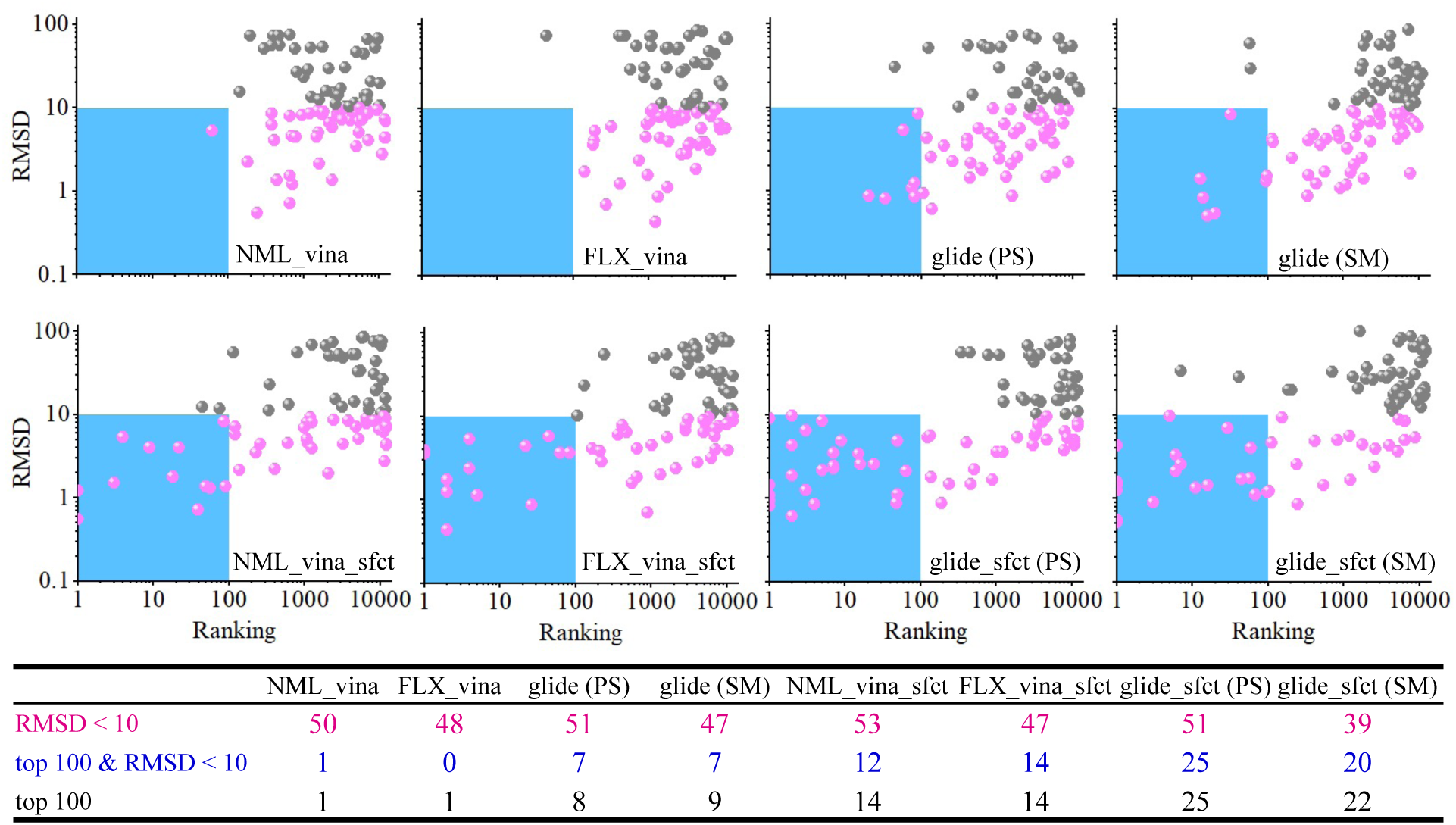
Reverse docking performance of the best pose of each test ligand and its original target. Each method was assessed based on its ability to correctly identify the binding pocket (RMSD < 10 Å) within the native target.

Among the top 100 predicted targets, there are occasional FPs. For example, FLX_vina has only one target ranked at 100, and it is considered FP (RMSD > 10 Å). The remaining pipelines also have one or two FPs. Nonetheless, the limited occurrence of false positive outcomes does not substantially influence the general pattern of results across the different pipeline approaches. However, the limited FPs did not affect the performance of these reverse docking processes. In our tests, Autodock Vina had almost no predictive ability, while Glide showed some predictive ability. After the SFCT correction, the performance of both approaches improved significantly, with glide_sfct performing the best. Although NML_vina_sfct had the highest number of successful reverse docking results, overall performance was better with glide_sfct (PS). This shows that both site prediction and scoring functions can impact reverse docking performance, but the superior ranking power of glide_sfct has a greater influence compared to site prediction.

### 2. The Impact of AF2 Structure Quality: Overall Accuracy and Side-chain Flexibility

To verify the effects of ginseng, Park et al. ^54^ performed reverse docking on a disease database with 1078 PDTD and kinase proteins. Li et al.^39^ selected 7769 protein-ligand complexes from the PDBbind database (version 2018) as the structure pool to evaluate the performance of 4 classical docking programs. In 2023, Trawally et al. ^55^ conducted reverse docking to understand the mechanism of action of a potent antimycobacterial compound 6g. The protein database consists of nine putative targets with fifteen different experimental structures. In these previous reverse docking studies, experimental structures were commonly used.

However, our work utilized structures predicted by AF2. Although AF2 generally achieves a confidence score of approximately 85 (plddt) for most protein structure predictions, indicating a significant level of accuracy in predicting overall protein structures, there is still room for improvement, particularly in predicting ligand binding pockets. For example, errors in predicting individual aromatic amino acid side chains during pocket prediction can lead to the closure of the entire pocket,^56,57^ which presents challenges for semi-flexible docking and can result in poor docking outcomes and potential false negative rates. In a study by Holcomb et al., AutoDock-GPU was used to redock AF2-predicted structures against experimental co-crystal structures.^58^ Their results revealed subpar docking performance of the predicted structures. In our results, macrophage metalloelastase (UniProt ID P39900) ^59^ has not been correctly identified by any pipelines (Figure S4). While the pocket prediction method, SiteMap, successfully predicted the binding site of the ligand, the AF2 structure has two additional structural domains compared to the experimental structure (Figure 7A). The ligand-binding site is occupied by a loop in the AF2 structure, preventing the successful docking of the small molecule at this site.

**Figure 7:**
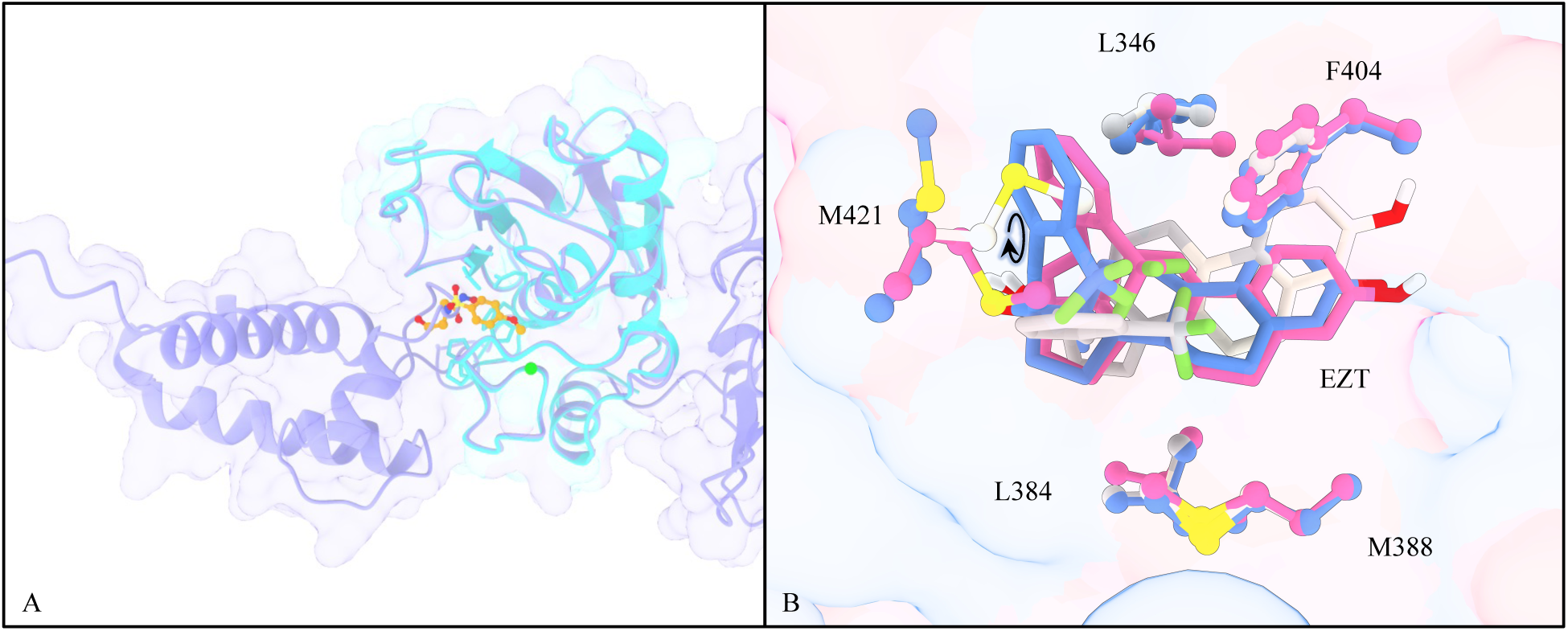
The quality performance of the structure predicted by AF2. (A) Structural comparison of the experimental and predicted structures of P39900. The experimental and AF2 predicted structures are colored in cyan and blue-violet, respectively. The ligand in the experimental complex is shown as balls and sticks, while the center of the predicted ligand-binding site is indicated by a green ball. (B) A study case of the effects of side-chain orientation of pocket residues on ligand binding: P03372 and EZT (PDB ID: 2p15). The protein is shown as cartoon, and ligands and key pocket residues are represented as sticks. The crystal structure is colored in cornflower blue, and the rigid and flexible docking poses generated by vina are colored in white and hot pink, respectively.

According to Figure 4D, FLX_vina_sfct is able to identify multiple targets that other pipelines fail to recognize. We speculate that this ability may be related to the flexibility of pocket side chains. For example, in the case of P03372, which has four ligands (WST, EST, JJ3, and EZT), NML_vina_deepRMSD, NML_vina_sfct, and FLX_vina_sfct pipelines successfully predict the first three ligands. However, for EZT, only flexible docking produced accurate predictions. In the FLX_vina_sfct, P03372 achieves the second highest rank, while in the NML_vina_sfct, it is ranked 11,518th. To investigate the reason, we compared the experimental structures and rigid and flexible docking poses (Figure 7B). After flexible docking, the amino acid residues that undergo changes were L346, L384, M388, F404, and M421. Among them, M421 exhibited the most significant variation. In the AF2 structure, the side chain of M421 occupied the position of the benzene ring in the experimental structure of EZT, causing the benzene ring of EZT to shift downward after rigid docking. However, the C*_α_*-C*_β_* bond of M421 rotated in flexible docking, allowing the ligand’s benzene ring to maintain its original orientation. This case demonstrates that flexible docking might serve as an important complement to improve hit rates when AF2 exhibits low pocket accuracy.

### 3. The Limits and Future Directions of Our Reverse Docking Pipeline

Previous reverse docking works utilize experimental protein structures, which ensured high structure quality and correct binding sites. However, the protein pool used had a limited number of protein targets that are not available at the human proteome level. Additionally, due to the limited ranking power of the existing scoring functions, the screening results often have a high rate of FPs. In our present work, we used human protein structures predicted by AF2 for reverse docking, which might rise the following concerns.

First, the binding sites are unknown and need to be predicted. The presence of multiple binding pockets and the failure to predict these pockets can lead to failures in reverse docking predictions. For example, PointSite failed to predict the binding sites of proteins O60885, P39086, P39900, and Q9H2K2. Consistently, all pipelines involving PointSite were unable to predict these targets. For SiteMap, it did not successfully predict the binding sites of the proteins O60674, P24941, and Q16539. Among them, the pipeline containing SiteMap successfully predicted the binding site of O60674, but failed to predict the binding sites of the remaining targets. After conducting a detailed examination of the results, we found that the protein O60674 was ranked among the top 58 hypothetical targets of JAK (PDB ID: 4f09) according to glide (SM). The RMSD between the experimental structure of JAK and the AF2 docking structure was 59.64. This indicates that the prediction of O60674 also failed, albeit being one of the two FPs in glide (SM). Another interesting case is that only SiteMap successfully predicted the binding site of bromodomain-containing protein 4 (UniProt ID O60885). ^60^ Therefore, even after SFCT correction, NML_vina_sfct, FLX_vina_sfct, and glide_sfct (PS), which predicted the binding site using PointSite, did not predict the target. However, glide_sfct (SM) successfully predicted the target. Therefore, if the site prediction fails, the overall prediction is unlikely to be successful.

Second, the side-chain quality of AF2-predicted structures might not be optimal, which can result in potential false negatives during docking. Flexible docking methods can predict targets that rigid docking cannot accurately predict, rigid docking methods can also predict targets that flexible docking cannot accurately predict. Although they complement each other in terms of predictive capabilities, there is no significant advantage in terms of the number of predictions made. Furthermore, it is unknown whether the proteins exist in an active or inactive conformation.

According to our findings, there are three considerations that may help address the aforementioned potential problems. Firstly, the AF2 human protein database is vast and contains structurally similar proteins. When a sizable set of proteins similar to the target of a particular ligand is available, they can fully showcase the commonalities and diversities of binding pockets in various proteins that interact with the compound, as well as the potential inclusion of both active and inactive conformations. Secondly, the test set consists of multiple structurally similar small molecules targeting the same protein. Similar small molecules have similar skeletons. Due to the appropriate redundancy of protein and small molecule structures, conformational sampling is enhanced, which helps us to discover a certain skeleton that can match multiple shared pockets in terms of geometry and energy. This promiscuity is the core of the problem. Lastly, the novel scoring approach that integrates a scoring correction such as SFCT provides more powerful and robust results.

Based on the work of Wang’s group, among the reported scoring functions, GlideScore-SP ranks second in terms of reverse screening capability, with ChemPLP@GOLD being only slightly higher than GlideScore-SP. ^21^ Li et al.^39^ evaluated the performance of 4 popular docking programs (AutoDock, Auto-Dock Vina, Glide and GOLD) in reverse docking. Their results show that Glide had the best capacity to find native targets. Thus, it is reasonable to speculate that glide_sfct (PS) is currently the most suitable pipeline for reverse docking tasks.

## Conclusion and Perspective

In the traditional drug development process, there is often a lack of a comprehensive evaluation of the interaction between drug small molecules and the human proteome. This can result in off-target effects and excessive toxicity when drugs are applied clinically, leading to failures. Reverse docking is an important tool to fill this gap. Here, we conducted an extensive reverse docking benchmark on AF2-predicted human proteins and 90 small molecules by utilizing two well-regarded docking programs, AutoDock Vina and Glide. By integrating the Vina, Gscore, SFCT, deepRMSD, and RTMscore scoring functions with two binding site prediction methods, we developed 11 different reverse-docking pipelines. Our results emphasize that each stage in reverse docking is vital, including the quality of protein structure and predicted binding site, and the evaluation of binding strength between ligands and receptors. Particularly, the combination of SFCT exhibits a higher ranking power than Gscore or Vina itself, two current state-of-art scoring functions. Our results also suggest that it is challenging to screen a specific target from thousands of proteins currently, but it is much easier to find proteins that have similar structures with the target. For example, in our work, the success rate of glide_sfct (PS) is only 27.8%, but when considering protein structures similar to the target, the success rate increases to 77.78%. Considering multiple similar small molecules to the query molecule for prediction, 92% of the targets can be successfully predicted. Therefore, in our future work, we can use the predictions of all human proteins as a starting point, and then filter the candidates for experimental evaluation by full-structure and pocket similarity. To summarize, our research revealed the performance of using AF2 for structure-based reverse docking to predict targets, and provided a highly useful pipeline for drug development. However, there are some limitations to this work. For example, the testing set primarily focuses on enzymes and receptors, considering no membrane proteins, such as GPCRs and ion channels, and the potential target database in the present work only considered single chain proteins and does not yet fully cover the human proteome by excluding some structures of poor quality.

## Acknowledgement

JG acknowledges the internal grant of Macao Polytechnic University (RP/CAI-01/2023), and YM acknowledges Singapore Ministry of Education (MOE), tier 1 grant RG97/22. The authors thank Dr. Lin Wang from ShanghaiTech University and Dr. Yinlei Han from Zhejiang University for their technical support.

## Data Availability Statement

All data can be obtained from https://github.com/molu851-luo/Reverse-docking-benchmark.git.

## Support Information

**Figure S1:**
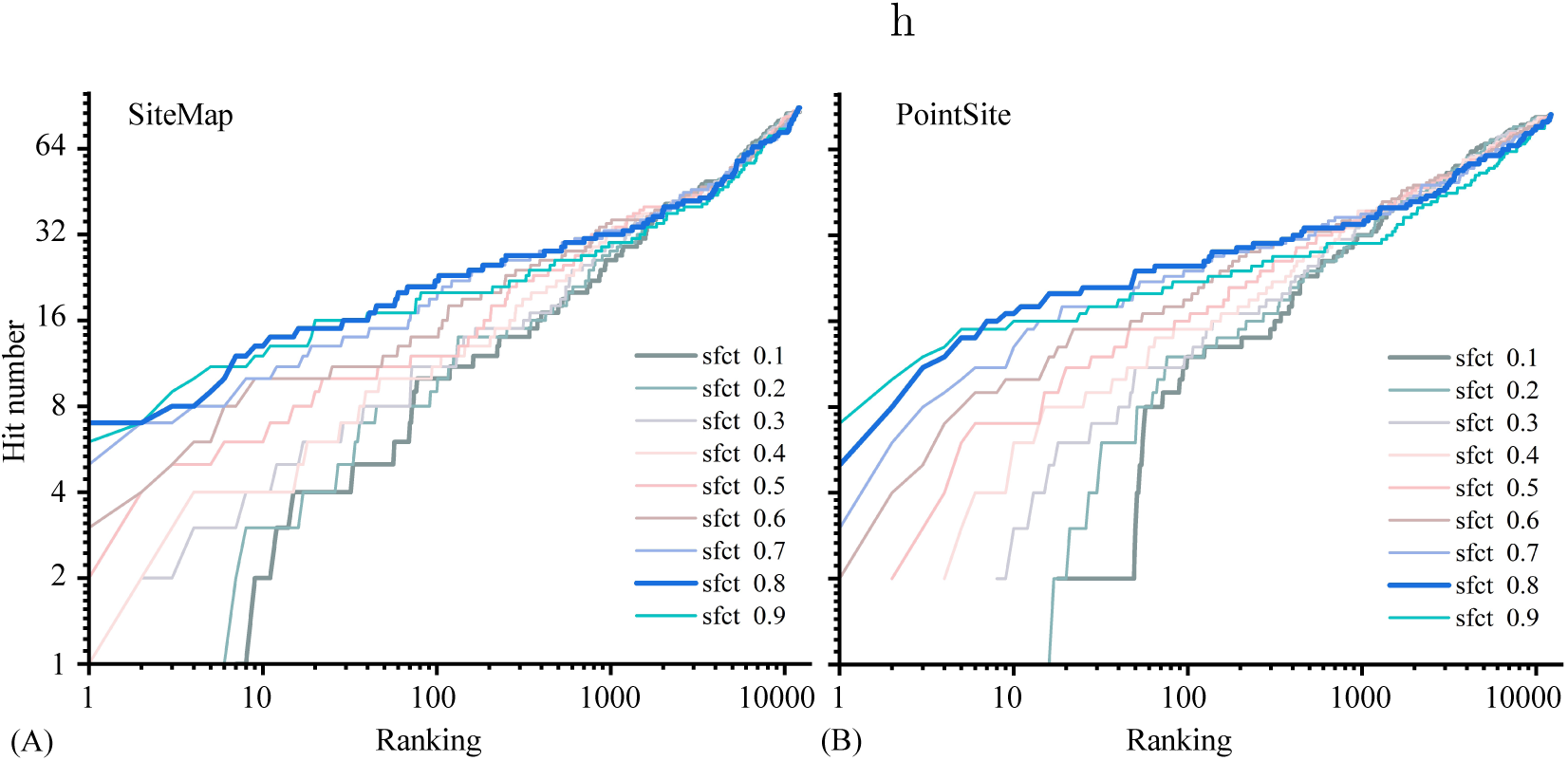
The performance of the combination of Glide+SiteMap and sfct.

**Figure S2:**
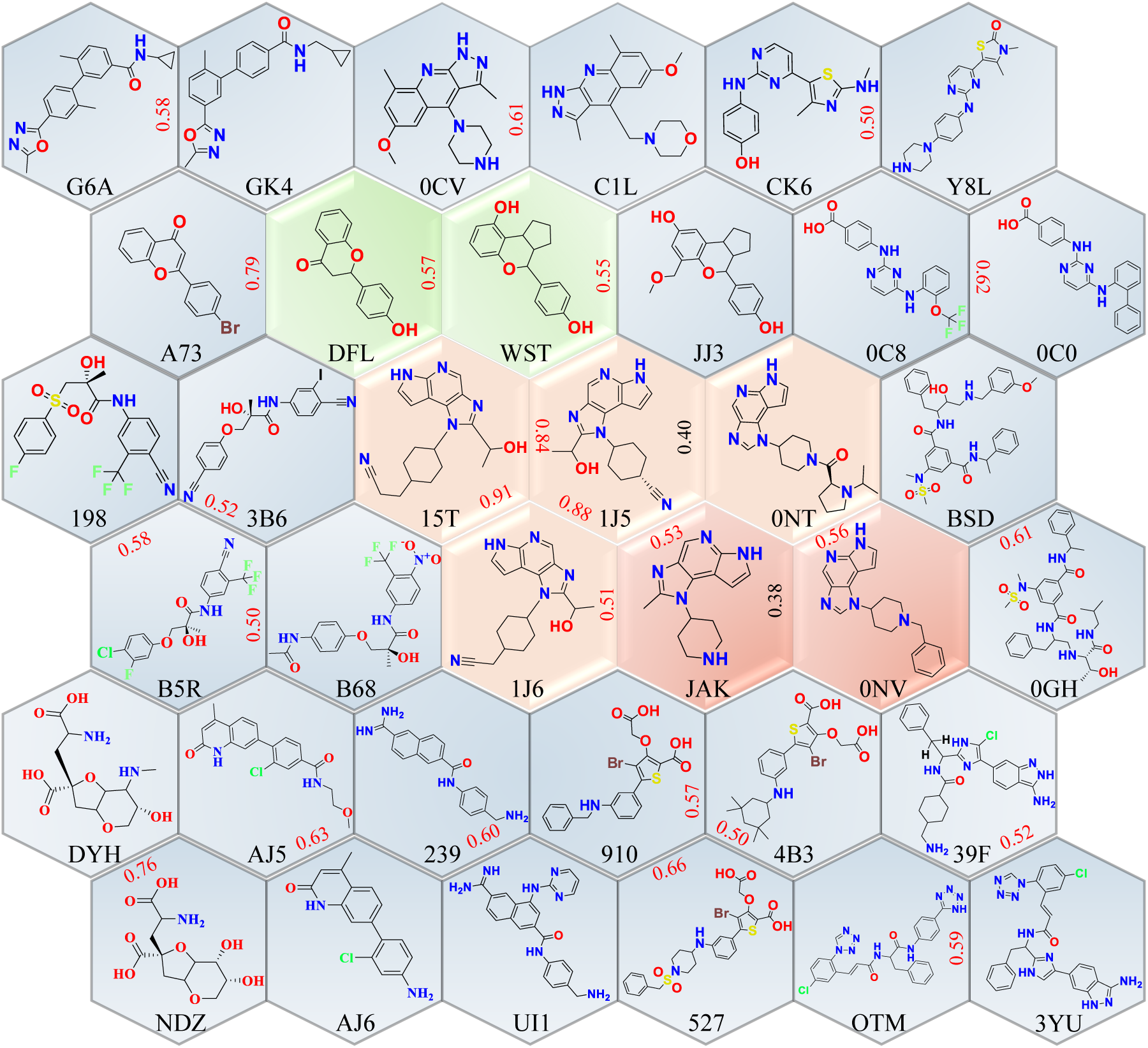
25 pairs of molecules with a similarity greater than 0.5. The deer color represents small molecules that are ligands for the target P23458. There are a total of 4 molecules in this category, and among them, 3 pairs have a similarity greater than 0.8. The sky blue represents pairs with the target O60674. P23458 and O60674 belong to the same protein family. The green indicates two similar small molecules, but their targets are unrelated. The remaining similar pairs of small molecules all target the same protein. The red numbers between each pair of molecules represent the similarity between them.

**Figure S3:**
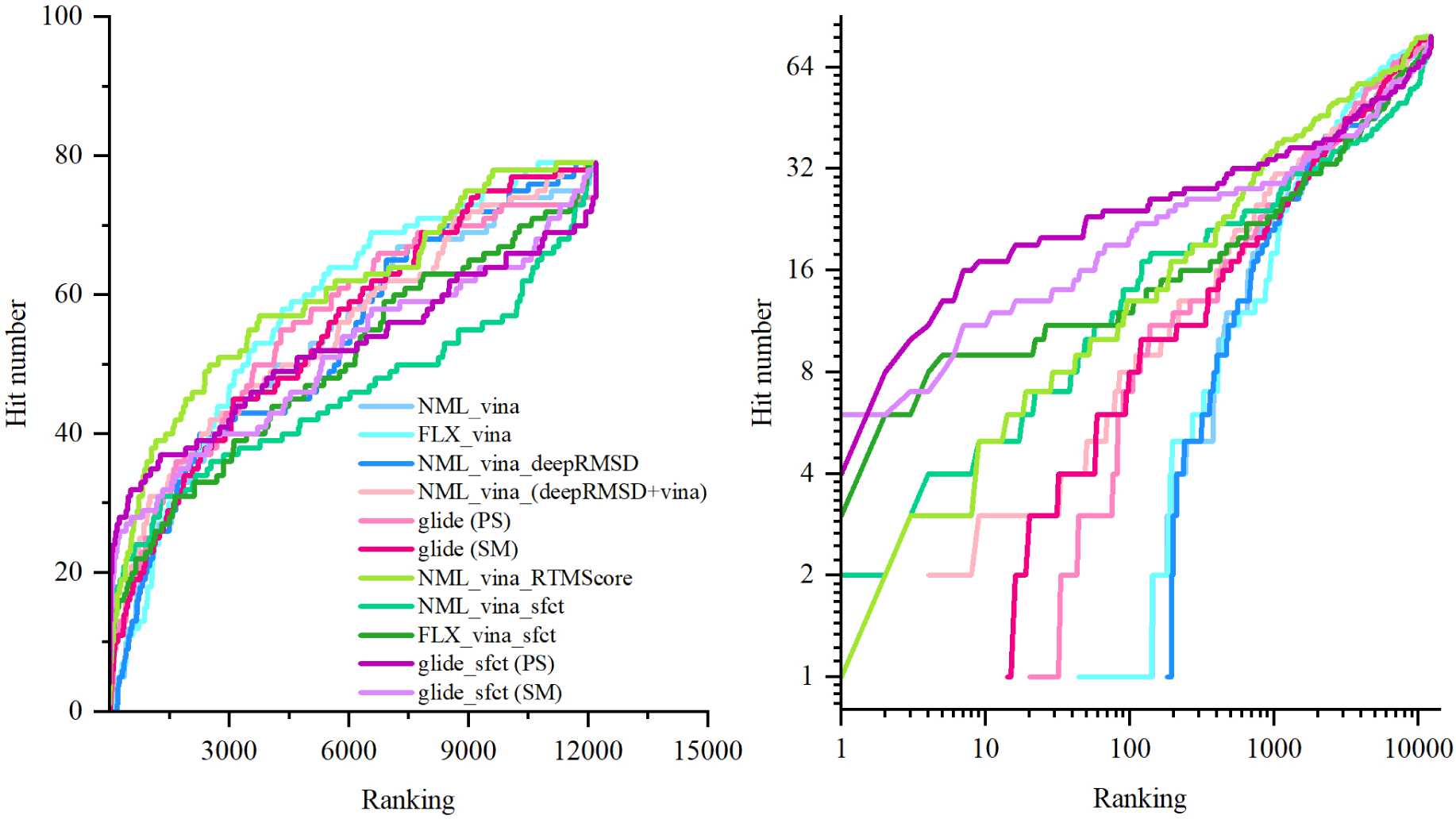
The performance assessment of distinct reverse-docking pipelines. (A–B) The plot of hit number as a function of ranking.

**Figure S4:**
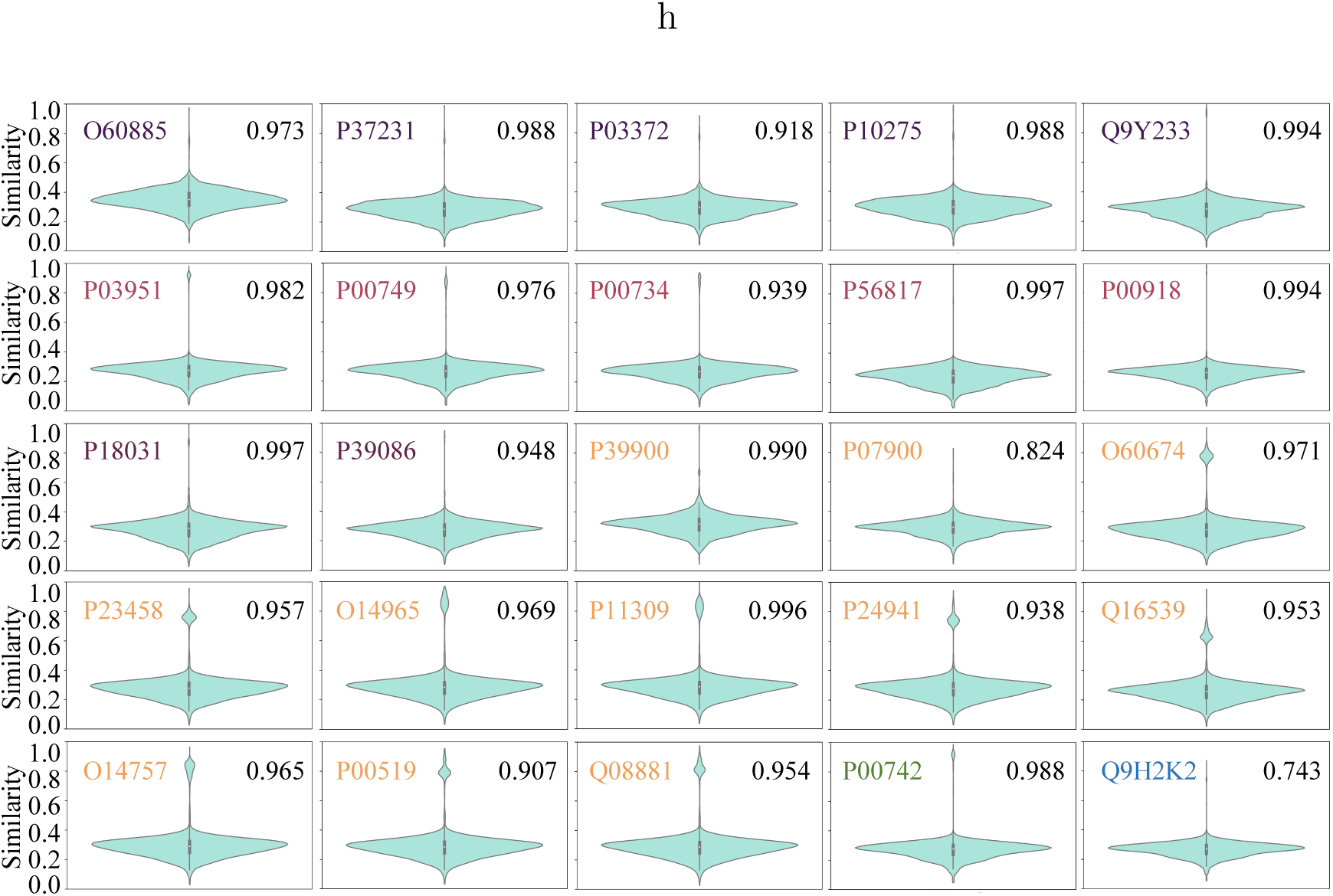
The distribution of the protein structure similarity (TM-score^41,61^ calculated by DeepAlign^40^) of each evaluated target and all processed human proteins.

**Figure S5:**
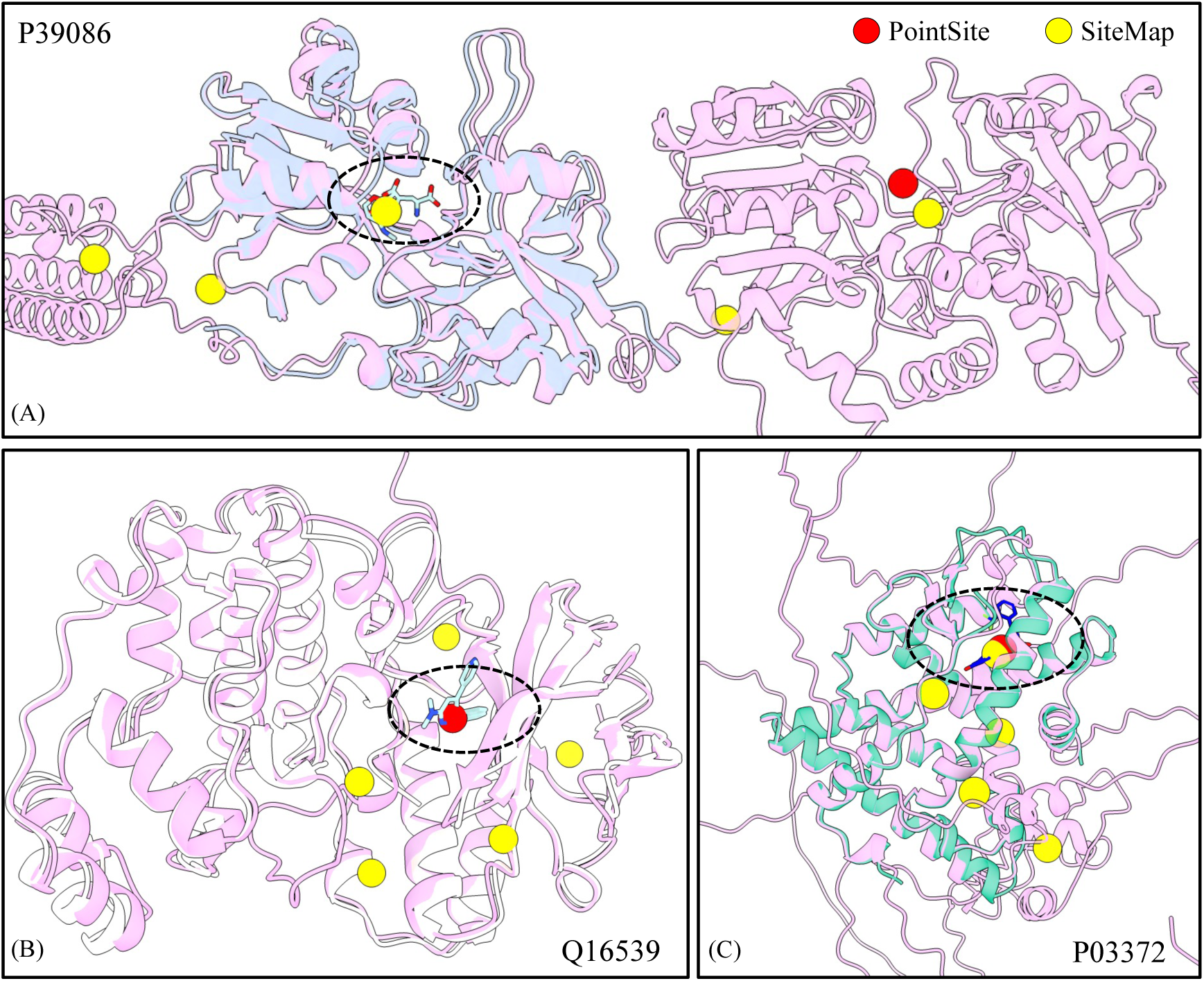
The study cases of binding site prediction: (A) P39086; (B) Q16539; (C) P03372. The AF2-predicted structures are colored in pink, and the experimental structures in blue, green, and white, respectively. Ligands in the experimental complexes are shown as sticks, and the sites predicted by PointSite (red) and SiteMap (yellow) are indicated by balls

**Figure S6:**
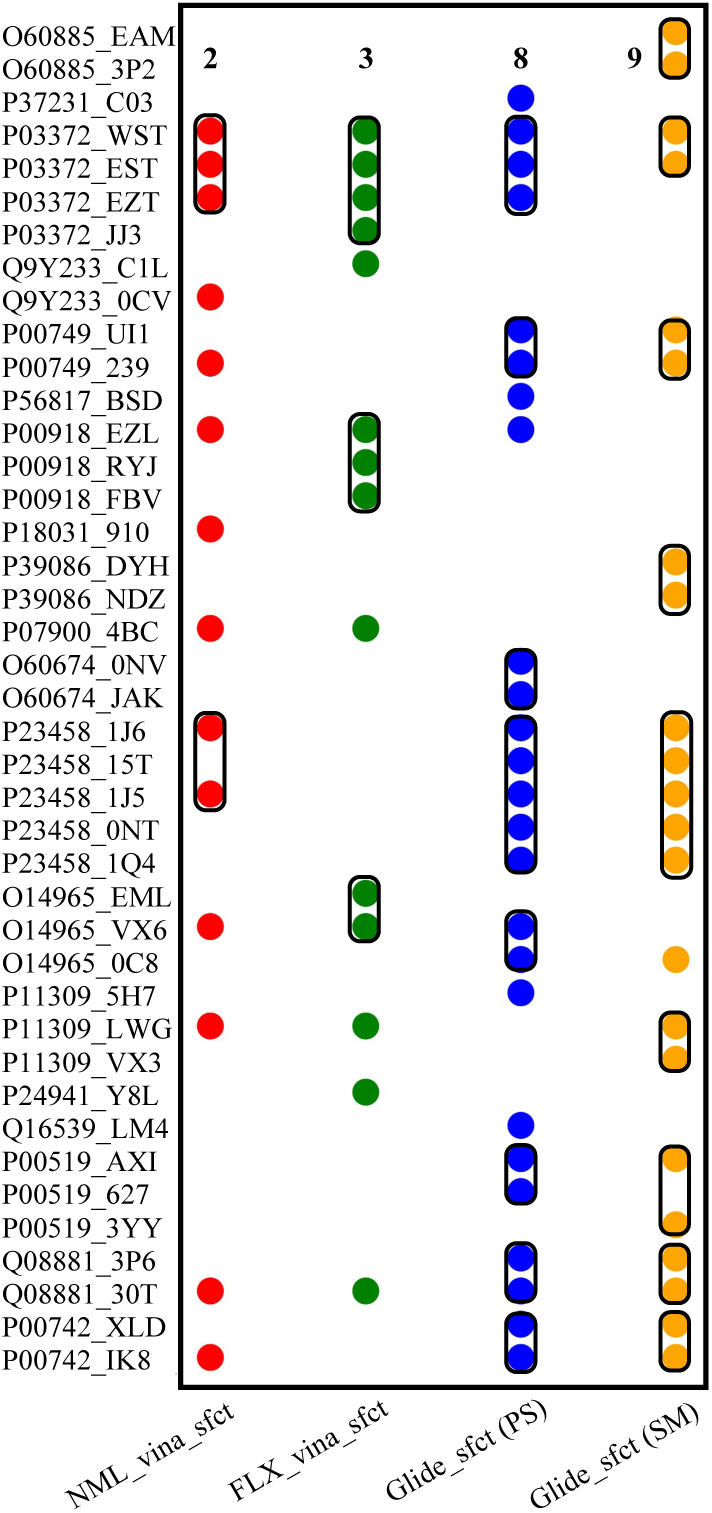
The four pipelines, which incorporate SFCT correction, successfully predicted ligands and targets. Within each box, the points represent small molecules that target the same protein

